# A systematic review quantifying host feeding patterns of *Culicoides* species responsible for pathogen transmission

**DOI:** 10.1101/2024.07.25.605155

**Authors:** Emma L Fairbanks, Michael J Tildesley, Janet M Daly

## Abstract

*Culicoides* biting midges are significant vectors of various pathogens, impacting both human and animal health globally. Understanding their host feeding patterns is crucial for deepening our understanding of disease transmission dynamics and developing effective control strategies. While several studies have identified the sources of blood meals in *Culicoides*, a quantitative synthesis of their host preferences and the factors influencing these behaviours is lacking. A systematic literature search focused on gathering data on (1) host selection and (2) host preference. For reviewing host selection we focused on studies reporting the identification of blood meal sources in individual *Culicoides*. When reviewing host preference we focused on studies comparing the number of *Culicoides* caught on or nearby different host species at the same location. Analysis revealed that some *Culicoides* species exhibit fixed host preferences, consistently feeding on specific hosts such as cattle and horses, while others display more opportunistic feeding behaviours. Notable variations were observed across different geographic regions. The findings indicate that host availability significantly influences *Culicoides* feeding patterns. This study highlights the complexity of host selection in *Culicoides* biting midges, which has implications for disease transmission. The variability in feeding behaviours underscores the need for regional assessments to inform targeted vector control strategies.

## 1 Background

*Culicoides* need a blood meal in order to develop eggs and produce offspring. While feeding on hosts they may act as vectors in the transmission of pathogens. Vector-borne diseases transmitted by *Culicoides* can severely impact human and animal health, as well as economies.

The World Health Organization (WHO) established four criteria for the recognition of a vector in the 1960s: (1) detection of the pathogen in unfed, field-collected insects; (2) demonstration of infection of the insect with the pathogen by feeding on an infectious host or artificial substitute; (3) demonstration of successful transmission of the pathogen from the insect to a susceptible host during feeding; (4) field evidence of association in space and time between the insects and susceptible hosts [1]. The fourth criterion is commonly explored by analysing the blood meals of field caught vectors; if the vectors are feeding on the susceptible hosts then this criterion is met.

Which hosts *Culicoides* feed on is influenced by availability (whether a host is present) and preference (what the vector prefers to feed on). For some vectors, availability has more of an influence on host selection, often referred to as opportunistic feeding. Alternatively, vectors more influenced by host preference are said to have fixed feeding patterns. These feeding behaviours are often explored by analysing the proportion of blood meals which originate from each host. For opportunistic vector species the proportion from each host type can be variable within small geographical regions, whereas the proportions are more consistent for vector species with fixed feeding behaviour.

Host selection can significantly impact disease transmission dynamics. If a vector that is capable of transmitting a pathogen regularly feeds on a species that is susceptible to infection with it, the capacity for disease transmission is much higher than if they rarely feed on that host species. It has long been understood that host selection plays a significant role in vectorial capacity, which estimates the number of new infectious cases that result from a vector feeding on a single infectious host on a single day [2]. Integral to this is the duration of the extrinsic incubation period (EIP), which refers to how long after feeding on an infectious host *Culicoides* become infectious, the probabilities of vector-to-host and host-to-vector transmission, the life expectancy of *Culicoides* and — of relevance here — the frequency with which the *Culicoides* species under consideration takes a blood meal from the host species. The proportion of blood meals from humans, referred to as the human blood index (HBI), is sometimes used to inform mathematical models for disease transmission [3–6].

While previous reviews have summarised the hosts identified in *Culicoides* blood meals [7, 8], there is a notable lack of quantitative reviews on host selection and the factors influencing feeding patterns. This study aims to fill this gap by systematically analysing host selection and host preferences of *Culicoides* species responsible for pathogen transmission. We conducted a comprehensive systematic literature search across multiple databases, focusing on studies that identified blood meal sources at the species level or compared the number of *Culicoides* attracted to different host types. Eligible studies were included based on predefined criteria to ensure robustness and relevance.

## 2 Methods

### 2.1 Systematic literature search

A systematic search (limited to the title and abstract) was performed (by E. L. Fairbanks) using PubMed, bioRvix, medRvix, arXiv, vectorbase, Paperity, Pubmed central and CAB direct on 19/09/2023. The search term was (“culicoides” OR “biting midges”) AND (“human blood index” OR “HBI” OR “host preference” OR “trophic preference” OR “meal preference” OR “blood preference” OR “blood meal” OR “blood-meal” OR “meal source” OR “blood source” OR “host blood” OR “anthropophilic index” OR “blood fed” OR “forage” OR “host use” OR “host associations”), adapted from a search term used to identify the HBI of Anopheles malaria vectors [9].

Duplicate studies were removed and the title and abstracts of the remaining articles were read and kept if they referenced the collection of *Culicoides* and were not review articles. The full text of articles which passed this criterion were read. Studies were eligible for data extraction for the blood meal review, to inform feeding ratios, if they identified the blood meals at the individual *Culicoides* level. Studies were eligible for data extraction for the host preference review if they compared the attractiveness of at least two host species by collection of *Culicoides* on or in close proximity to the hosts in the field.

### 2.2 Data extraction and analysis

Studies that reported *C. pallidipennis* were recorded as *C. imicola*, due to the renaming of this vector. Species within the Obsoletus and Variipennis complexes were pooled for analysis because studies which use morphology for species identification would not be able to determine the specific species. The Variipennis complex consists of five subspecies: *C. variipennis*, *C. sonorensis*, *C. occidentalis*, *C. australis* and *C. albertensis*. The Obsoletus complex consists of three species: *C. obsoletus*, *C. scoticus* and *C. montanus*. Other members of the Obsoletus group, *C. chiopterus*and *C. dewulfi*, can be identified morphologically and were therefore analysed separately.

Data analysis was performed using Rstudio [10]. Figures were generated using the *ggplot2* package in Rstudio [10, 11].

#### 2.2.1 Blood meal review

For the quantitative blood-meal review, data extracted from each article included: country, location (including latitude and longitude where reported), years sampling took place, trapping methods, *Culicoides* species identification methods, blood meal identification method and, for each *Culicoides* species or complex, the number of identified blood meals and how many where attributed to each host type.

Some studies included collections at more than one location; data from these were extracted separately. If the results reported were for pooled locations these were recorded as one data set with the mid-point between the locations as the latitude and longitude. If a blood meal was identified as from a particular host species at a location, it was assumed to be present at that location. Therefore, *Culicoides*species that were not reported to have fed on the host at the location were recorded as having 0% of the blood meal coming from the host type. This was included in calculations of the weighted mean, weighted 95% confidence interval and range of the percentage of blood meals from each host type that is susceptible to a disease for which the *Culicoides* species/complex is a competent vector. Sheep and goats were not considered to be susceptible to vesicular stomatitis virus [12].

These summary statistics were calculated for all host species and *Culicoides* species/complex combinations for which a pathogen was identified as infecting both the host and *Culicoides* species and evidence was found of the *Culicoides* species feeding on the host (the host was identified as a blood meal in at least one data point). Results were aggregated and summarised by *Culicoides* species/complex, host species and region (Americas, Asia, Africa, Australia or Europe).

The mean was weighted by the total number of the *Culicoides* species caught for each data point. To estimate the uncertainty associated with the observed proportions of host species in vector blood meals, bootstrap confidence intervals were calculated. This non-parametric approach involved resampling the original data with replacement to create numerous simulated samples. The process was repeated 1,000 times for each combination of vector species, host species and region.

For each bootstrap sample, the proportion of blood meals derived from a specific host relative to the total number of meals analysed was calculated. This step generated a distribution of mean percentages for each host species, from which the 2.5*^th^* and 97.5*^th^* percentiles were determined. These percentiles represent the lower and upper bounds of the 95% confidence interval, respectively, providing an estimate of the range within which the true mean percentage likely falls.

#### 2.2.2 Host preference review

For the quantitative host-preference review, data extracted included: country, co-ordinates (latitude and longitude, if available), years of collection, trapping methods, *Culicoides* species identification methods, *Culicoides* species, host attractant and the number of the *Culicoides* species caught on the host attractant.

For each study we compared the ratio of the number of each *Culicoides* species caught on each combination of host pairs. Where more than one study investigated a *Culicoides* species preferences for a host pair the mean and range of these ratios was calculated. These results were then compared between host preference studies and to results from the vector blood meal review.

If trapping was performed for more than one distance from the host, data were extracted for the closest distance to the host. This is because as the distance from hosts increases, the number of *Culicoides* caught decreases [13]. The closest location to the host is assumed to most accurately reflect the number biting the host.

## 3 Results

A total of 532 articles were found during the systematic search from Pubmed, bioRvix, Paperity, Pubmed central and CAB direct. Of these 141 were duplicates. No articles were found on medRvix, arXiv or vectorbase. Of the 391 unique articles, 308 were excluded during a first screening of their titles for being reviews or not including collection of *Culicoides*. After reading the full texts of the remaining 78 articles which could be retrieved a further 23 were removed for not collecting *Culicoides* or reporting the number of *Culicoides* found to have fed on each host species. During the screening, five titles were identified that qualified for a full text read but could not be accessed. In total, data were extracted from 55 articles; 45 analysed in the blood meal review and 11 analysed in the host preference review with one article containing data used in both reviews. The process of the literature search is described by the PRISMA flow diagram [14] in Figure 1.

**Figure 1:**
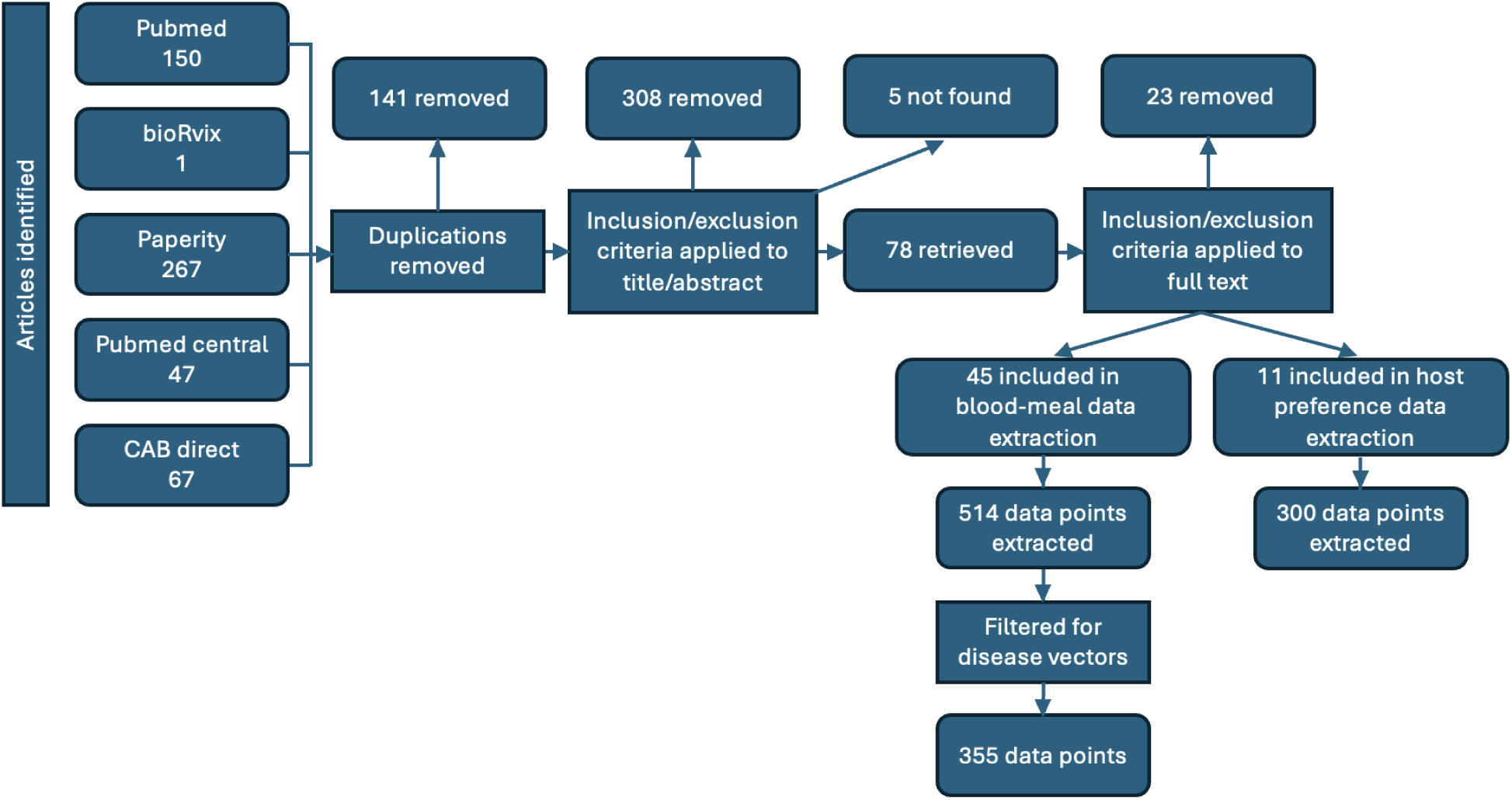
PRISMA flow diagram describing the process of the systematic search.

### 3.1 Blood meal review

All data generated from the blood meal systematic search are provided in Supplementary file 1. Overall, 514 data points were extracted from 45 studies including 124 locations in 26 countries [15–59]. Of these studies, 53% only gathered data from one location Figure 2. The most common trapping method was light-baited suction traps, used at 77% of locations. Data inspection revealed no significant differences in blood meal composition across trapping methods, suggesting that the choice of trapping technique did not substantially bias the results. Polymerase chain reaction (PCR) was used to identify the host from which blood meals where taken in 87% of studies and was used in all studies conducted and reported after 1992. Alternative methods for identifying blood meals included enzyme-linked immunosorbent assay (ELISA) in 4% and precipitin tests in 9% of studies. Of the 45 studies, 24 used morphology alone to identify *Culicoides* species, whereas 16 also used PCR and 1 also used PCR and mass spectrometry. Four studies did not specify how the species of *Culicoides* was identified. The most common geographical area analysed in studies was Europe (n=19), followed by Africa (n=8), the Americas (n=8), Asia (n=8) and Australia (n=2). There has been a rapid and substantial increase in the number of studies reported since 2007, especially in Europe, the Americas and Asia, possibly due to rising concerns around *Culicoides*-borne pathogens.

**Figure 2:**
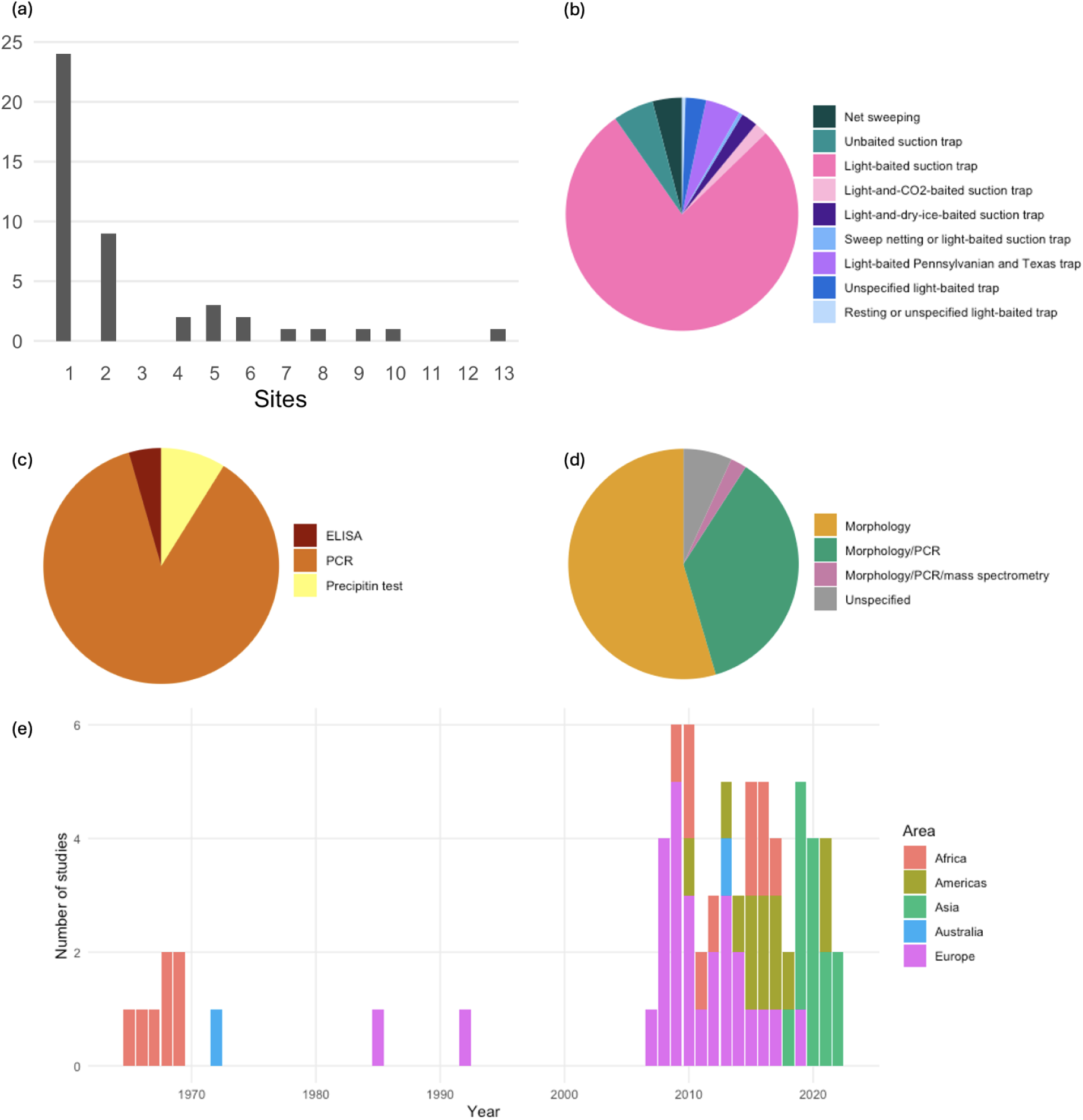
Summary of data collected from 45 studies for the blood meal review. (a) The number of sites in each study, (b) The method of catching *Culicoides* at each site, (c) the method of identifying the host source of blood meals, (d) the method of identifying *Culicoides* species and (e) the number of studies capturing *Culicoides* each year by geographical region.

After filtering for disease vectors that fed on competent hosts, 355 data points were analysed. The pathogens and corresponding vectors found to have fed on each host are listed by host species in Table 1.

**Table 1:**
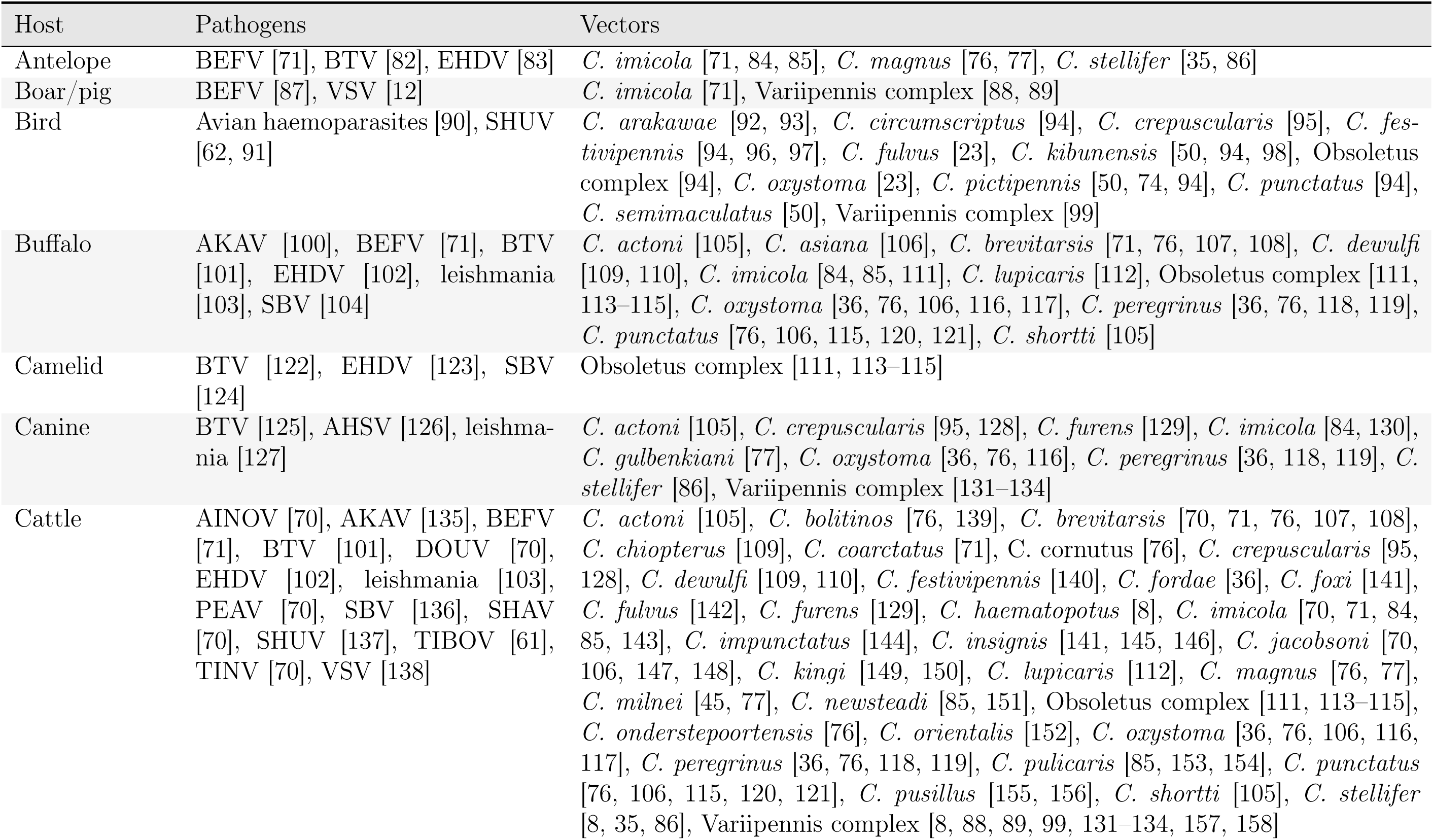
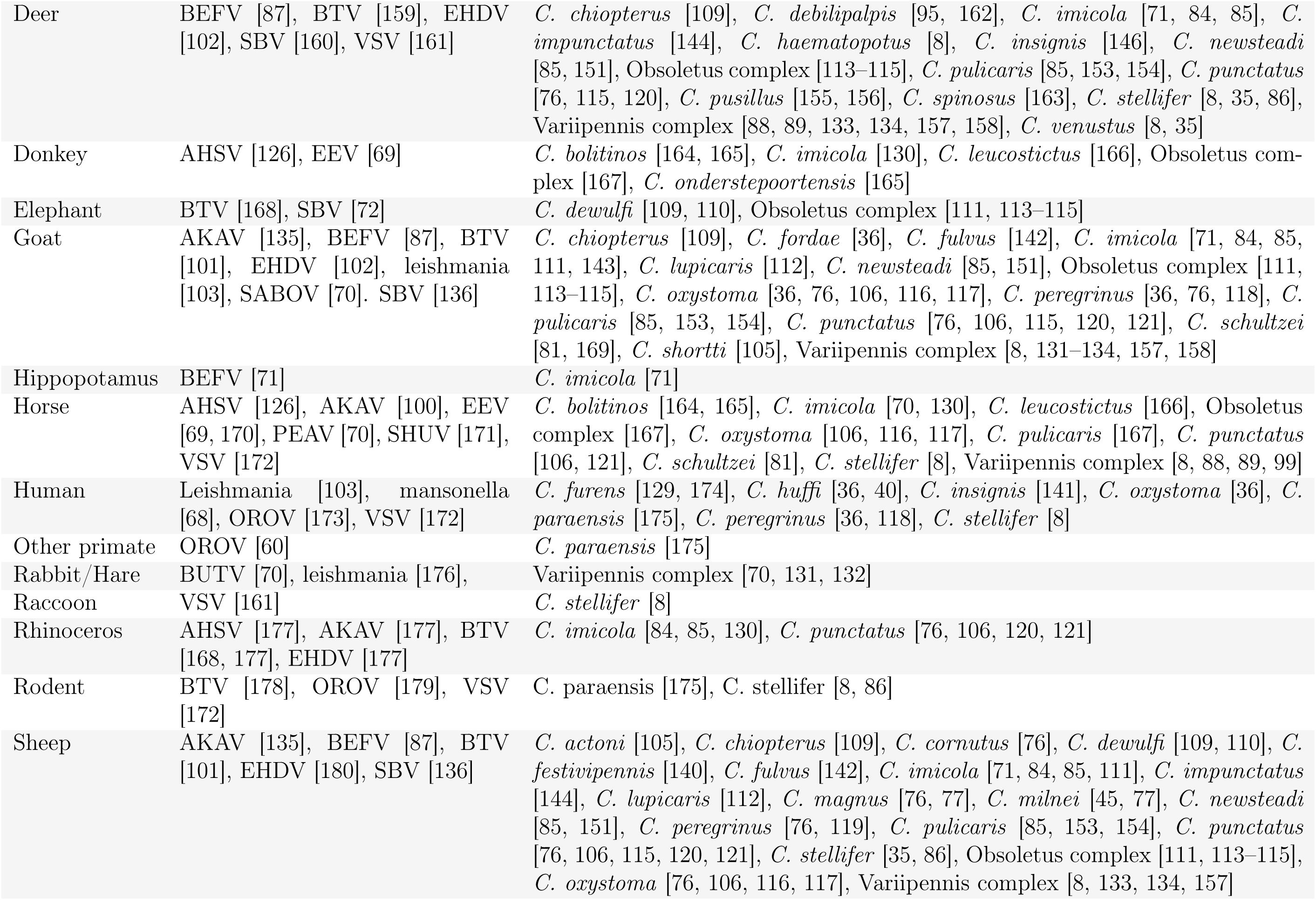
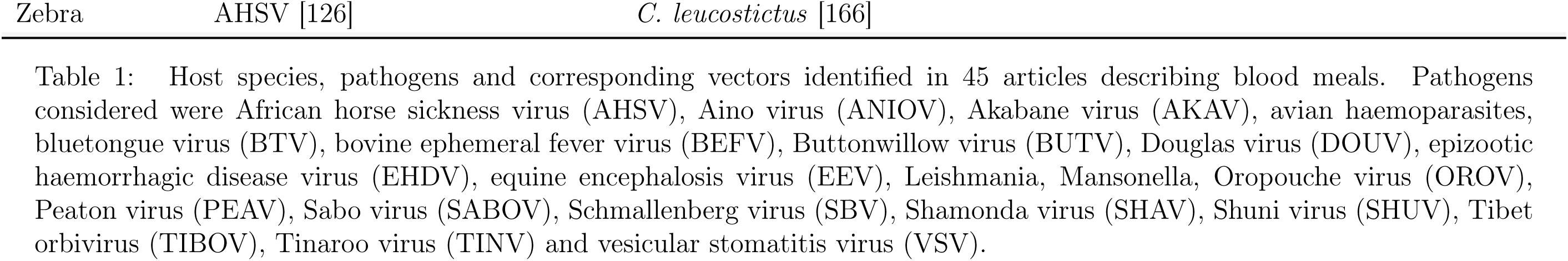
Host species, pathogens and corresponding vectors identified in 45 articles describing blood meals. Pathogens considered were African horse sickness virus (AHSV), Aino virus (ANIOV), Akabane virus (AKAV), avian haemoparasites, bluetongue virus (BTV), bovine ephemeral fever virus (BEFV), Buttonwillow virus (BUTV), Douglas virus (DOUV), epizootic haemorrhagic disease virus (EHDV), equine encephalosis virus (EEV), Leishmania, Mansonella, Oropouche virus (OROV), Peaton virus (PEAV), Sabo virus (SABOV), Schmallenberg virus (SBV), Shamonda virus (SHAV), Shuni virus (SHUV), Tibet orbivirus (TIBOV), Tinaroo virus (TINV) and vesicular stomatitis virus (VSV).

There are some pathogens to which vertebrate hosts listed in 1 are known to be susceptible, but these are not included in Table 1. Pathogens are only ascribed to a vertebrate host if there was evidence that a *Culicoides* species/complex associated with transmission of the pathogen fed on the susceptible host in the identified blood meals. For example, birds and sloths are known to be susceptible to Oropouche virus [60], goats and buffalo to Tibet orbivirus [61], buffalo, antelope, alpaca (camelid), giraffe, crocodile, rhinoceros and rodents to Shuni virus [62, 63], rodents, sloths, non-human primates, marsupials, cats, anteaters, shuckand raccoons to Leishmania [64–67], rodents and non-human primates to mansonella [68], zebra to equine encephalosis virus [69], sheep and goats for Aino virus [70], elephants, giraffe, swine and camelids to bovine ephemeral fever [71], giraffe to Schmallenberg virus [72], birds to Thimiri virus, as well as camelids to Akabane virus [73] and vesicular stomatitis virus [12].

Some *Culicoides* species have also been associated with transmission of pathogens for which there was no evidence that they had fed on susceptible hosts in the identified blood meals. For example, *C. pallidicornis*, *C. poperinghensis* and *C. impunctatus* for avian haemoparasites [50, 74, 75], *C. paraensis*, *C. zuluensis* and *C. nivosus* for epizootic haemorragic disease virus [76], *C. pycnostictus* and *C. bedfordi* for bluetongue virus [77], *C. leucostictus* for blue-tongue virus and epizootic haemorrhagic disease virus [76, 77], *C. endermania* for bluetongue virus, epizootic haemorrhagic disease virus and equine encephalosis virus [78, 79], *C. mahasarakhamense* for Leishmania [40], *C. austeni*, *C. grahamii* and *C. milnei* for mansonella [80] and *C. circumscriptus* for Akabane virus [81].

Table 2 presents the total number of the *Culicoides* species caught and the weighted mean, weighted 95% confidence interval and range of the blood index for susceptible hosts which they were found to have taken a blood meal from (Table 1) with data for at least 4 locations for each region. These summary statistics are provided for all *Culicoides* species and corresponding susceptible hosts in Table 1 for each region in Supplementary file 2.

**Table 2:**
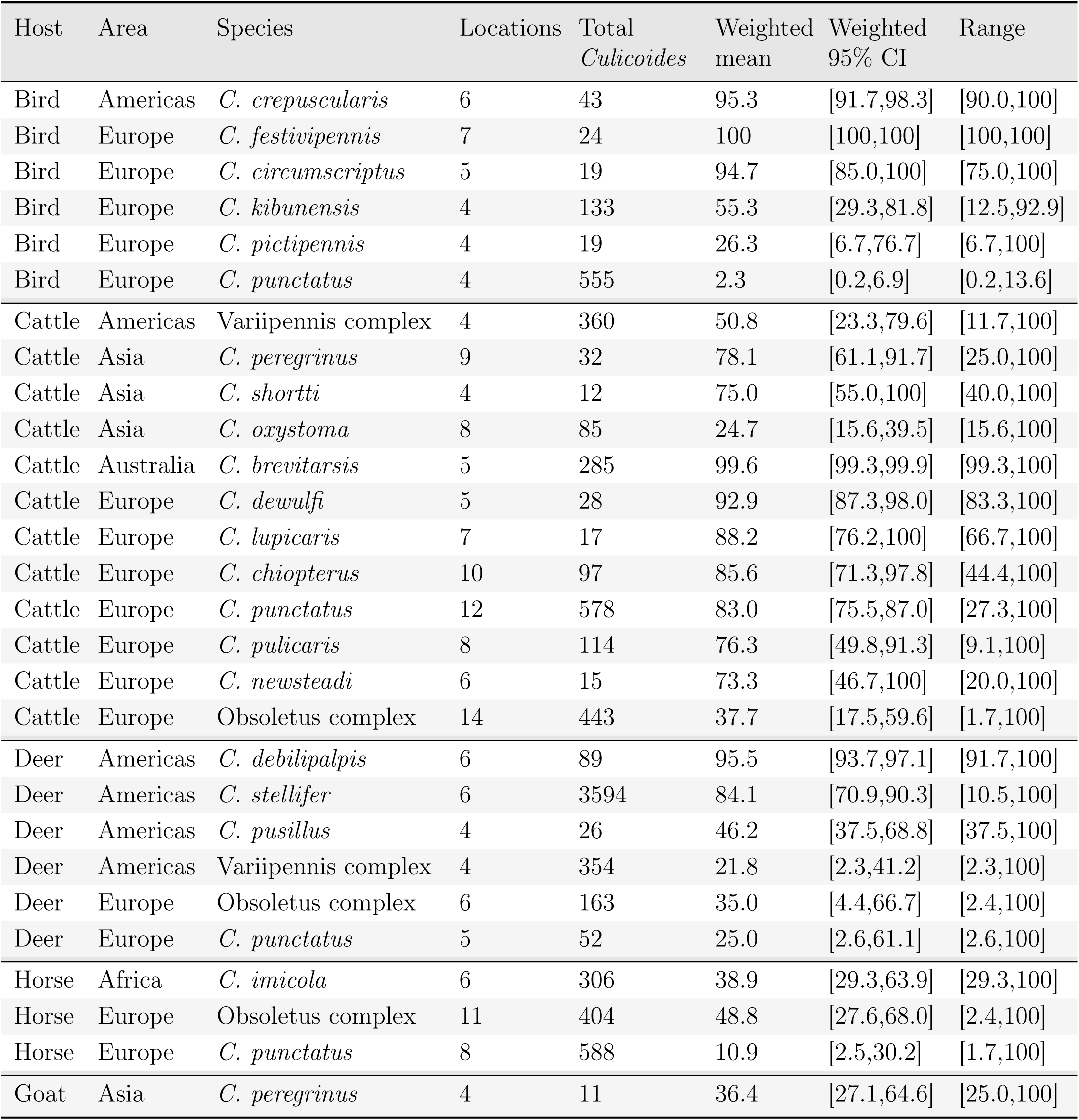
The total number of each *Culicoides* species with blood meals identified, the number of locations they were collected and the weighted mean, weighted 95% confidence interval and range of the percentage of blood meals from each susceptible host they were found to feed on given the *Culicoides* species was caught in at least 4 locations in a region.

Bluetongue virus has been recovered from wild rodents, however their role as virus reservoirs and potential implication in transmission is unclear [178]. Six studies reported *Culicoides* species susceptible to bluetongue virus feeding on rodents in the Americas and Europe (*C. crepuscularis*, *C. stellifer*, Obsoletus complex, *C. punctatus* and *C. debilipalpis*). Considering studies with more than four of the *Culicoides* species with blood meals identified, we found that less than 3% of these vector species had fed on rodents. This is lower than most susceptible host species, including antelope, buffalo, camelids, cattle, deer, elephants, goats, rhinoceros and sheep. Canids were fed on by competent bluetongue virus *Culicoides*species in nine locations, however similarly to rodents, the percentage of blood meals belonging to this host species is relatively low.

*Culicoides* species exhibit diverse host preferences, ranging from highly specific to opportunistic feeding behaviours. Some species demonstrate a strong preference for birds, such as *C. crepuscularis*, *C. festivipennis*, and *C. circumscriptus*, with weighted means for the percentage of blood meals taken from birds *≥* 95%. Conversely, other species show a clear mammalian preference, exemplified by the Variipennis complex and *C. punctatus*, with weighted means for the percentage of blood meals from non-mammalian hosts *≤* 5%. However, several species display more opportunistic host selection behaviours. The Obsoletus complex, *C. kibunensis*, and *C. pictipennis*, for instance, exhibit a broad range in the percentage of blood meals from birds across different locations, indicating flexibility in host choice.

Within mammalian-feeding species, further preferences are observed. Some species show a fixed feeding behaviour for specific mammals, such as *C. debilipalpis*, with weighted mean and range for the percentage of blood meals from deer of 95.5% and 91.7–100%, respectively. In contrast, other species demonstrate more opportunistic feeding within the mammalian class. Members of the Variipennis and Obsoletus complexes, for example, have been observed feeding on a variety of mammals including cattle, deer, and horses, suggesting a broader host range within mammals.

While dogs have been reported to become infected with African horse sickness virus [126], their role in the transmission cycle may be more complex. It is possible that dog infections occur through non-vector routes, such as ingestion of infected meat from horse carcasses [126]. We identified two studies which reported African horse sickness virus in the competent *Culicoides* vector species *C. imicola* and *C. oxystoma*, with 1% and 9% having fed on canines, respectively. This is substantially lower than percentages of blood meals belonging to horses for these species.

### 3.2 Host preference review

Overall, 252 data points were extracted from 11 studies including locations in 8 countries [13, 57, 181–189]. Each study only collected data from one location (Figure 3). *Culicoides* collection methods included box traps (such as drop traps or stables used to trap insects for collection by aspiration), aspiration directly from host species, sticky traps attached to host species and net sweeping close to host species. Of the 11 studies, 8 used morphology alone to identify *Culicoides* species, whereas 2 also used PCR and 1 did not specify the method. The most common geographical regions analysed in studies was Europe (n=7), followed by Africa (n=1), the Americas (n=1), Asia (n=1) and Australia (n=1). Most studies were performed in the Netherlands (n=4). Like the blood meal analysis there has been a substantial increase in the number of studies in recent years.

**Figure 3:**
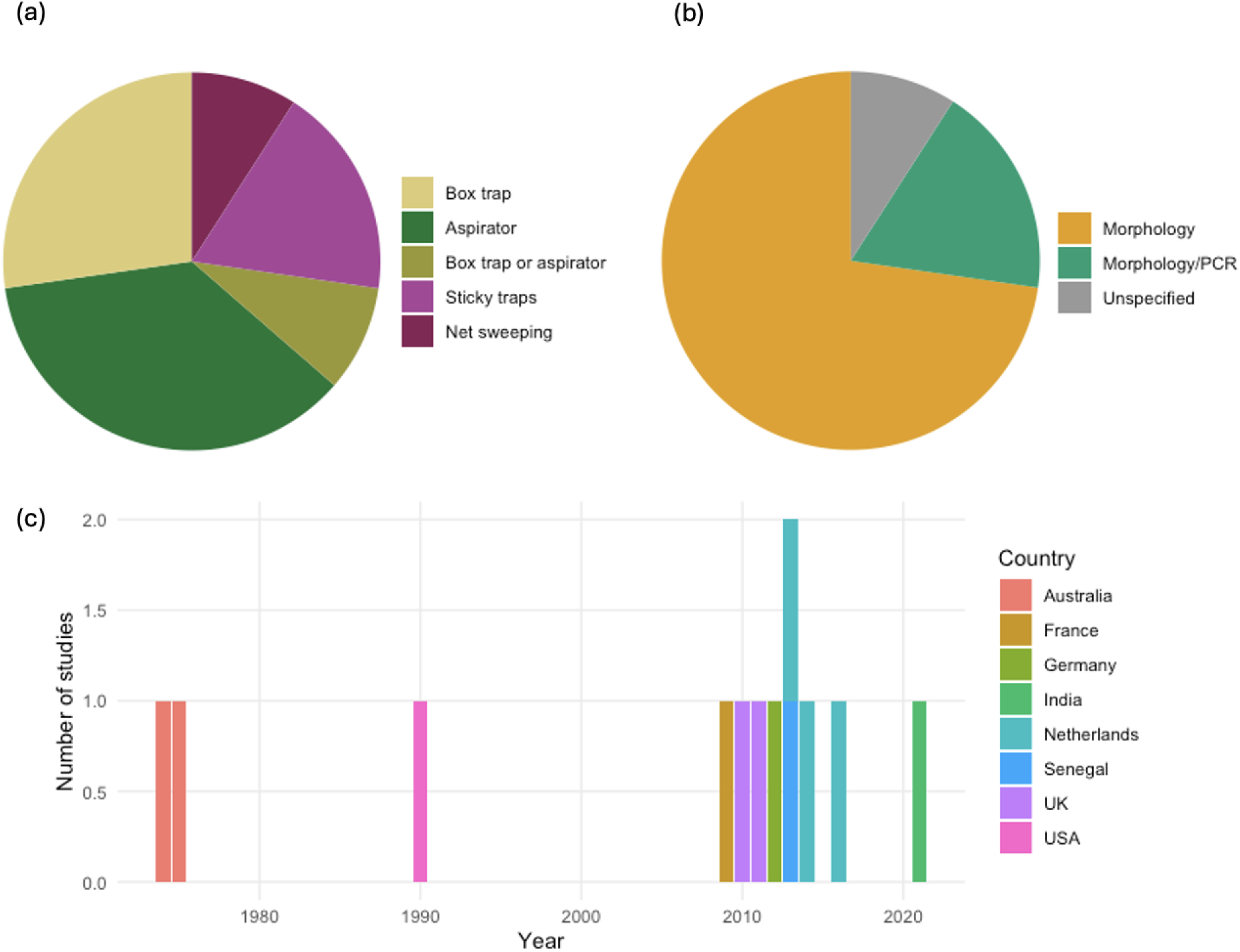
Summary of data collected from 11 studies for the host preference review. (a) The method of catching *Culicoides* in each study, (b) the method of identifying *Culicoides* species and (c) the number of studies capturing *Culicoides* each year by country.

Table 3 shows the results from host-preference studies for host pairs with at least two studies comparing their preferences for a *Culicoides* species. This result is provided for all host pairs investigated for each *Culicoides* species in Supplementary file 3.

**Table 3:** The mean and range of ratios of the number of vectors caught for host attractants for *Culicoides* species and host pairs investigated in at least 2 host preference studies. Studies marked with a *†* show the same study site was used for both studies. All data summarised in this table was from Europe.

| Host pair | Species | Studies | Total *Culicoides* | Mean ratio | Ratio range |
| --- | --- | --- | --- | --- | --- |
| Cattle:Horse | *C. fascipennis* | 2 † | 102 | 15.12 | [1.21,29.00] |
| Cattle:Horse | *C. chiopterus* | 2 | 4883 | 5.75 | [5.65,5.85] |
| Cattle:Horse | *C. pallidicornis* | 2 † | 204 | 4.83 | [4.82,4.83] |
| Cattle:Horse | *C. achrayi* | 2 † | 1855 | 2.44 | [1.79,3.10] |
| Cattle:Horse | *C. impunctatus* | 2 † | 20 | 2.38 | [2.25,2.50] |
| Cattle:Horse | *C. pulicaris* | 2 † | 429 | 1.96 | [0.49,3.44] |
| Cattle:Horse | *C. stigma* | 2 † | 237 | 1.80 | [0.85,2.75] |
| Cattle:Horse | *C. punctatus* | 2 † | 3841 | 1.74 | [0.88,2.59] |
| Cattle:Horse | Obsoletus complex | 3 | 7398 | 0.96 | [0.02, 1.66] |
| Cattle:Horse | *C. dewulfi* | 3 | 5451 | 0.63 | [0.01,1.18] |
| Cattle:Horse | *C. heliophilus* | 2 † | 36 | 0.48 | [0.14,0.82] |
| Cattle:Sheep | *C. achrayi* | 4 | 984 | 42.05 | [1.32,100.5] |
| Cattle:Sheep | *C. pulicaris* | 3 | 666 | 30.01 | [12.53,55.00] |
| Cattle:Sheep | *C. punctatus* | 4 | 3792 | 22.55 | [1.68,50.65] |
| Cattle:Sheep | *C. chiopterus* | 4 | 7566 | 14.14 | [1.24,24.21] |
| Cattle:Sheep | *C. dewulfi* | 5 | 8976 | 5.91 | [0.33,13.66] |
| Cattle:Sheep | Obsoletus complex | 5 | 15509 | 5.30 | [2.40,11.43] |
| Cattle:Sheep | *C. impunctatus* | 3 | 82 | 2.79 | [1.25,3.75] |
| Cattle:Sheep | Obsoletus group | 2 | 5471 | 1.03 | [1.00,1.06] |
| Horse:Sheep | Obsoletus complex | 2 | 2304 | 91.31 | [2.13,180.50] |
| Horse:Sheep | *C. dewulfi* | 2 | 2141 | 36.79 | [6.91,66.67] |

The mean ratios of the numbers of *Culicoides* caught on two host species varied from 0.48–91. Our analysis reveals varying degrees of consistency in host preference ratios across studies. For instance, the cattle-to-horse preference ratios for *C. chiopterus*, *C. impunctatus*, and *C. pallidicornis* show relatively low variability (ranges of 5.65-5.85, 2.25-2.50, and 4.82-4.83, respectively). In contrast, other species exhibit markedly higher variability in host preferences; for example, the cattle-to-sheep ratio for *C. achrayi* ranges from 1.32 to 100.5 across different studies. The level of variability does not appear to be associated with the location. When *Culicoides* species that were investigated multiple times at the same location, some studies found similar catch ratios between the hosts and others did not.

In this review, two studies reported results for all subspecies of the Obsoletus group pooled. These studies are reported separately to the Obsoletus complex. We observe that results for comparing cattle to sheep (Table 3) are not consistent between the Obsoletus group and Obsoletus complex, with the Obsoletus group showing less preference for cattle. This might be explained by the wide range of results found for *C. dewulfi*, suggesting it has very opportunistic feeding behaviours.

Table 4 shows the summary statistics for host pairs and corresponding *Culicoides* species with more than four data points in the blood meal review (included in Table 2). We observe that the trends from the host preference review generally agree with the blood meal review, in terms of the host most commonly selected. For *C. punctatus* the blood meal review found the mean percentage of blood meals belonging to cattle and horses was 83.0% (95% CI: [75.5,87.0]) and 10.9% (95% CI: [2.5,30.2]), respectively. However, the host preference review did not find evidence of a preference for cattle with the ratio of the number of vectors found on cattle compared to horses ranging from 0.88 to 2.59.

**Table 4:** Comparison of host selection ratios between host preference studies and blood meal analyses for *Culicoides* species. For the host preference studies the mean ratio of the number of vectors caught on different host attractants is given. For the blood meal studies the ratio of the means of the percentage of blood meals from each host type is given. Only comparisons with *≥* 4 blood meal data points per host are included.

| Host pair | Species | Region | Host prefer-<br>ence review | Blood meal review |  |  |
| --- | --- | --- | --- | --- | --- | --- |
|  |  |  | Mean ratio | Mean %<br>host 1 | Mean %<br>host 2 | Ratio |
| Cattle:Goat | *C. peregrinus* | Asia | 43.86 | 78.1 | 36.4 | 2.15 |
| Cattle:Horse | *C. punctatus* | Europe | 1.74 | 83.0 | 10.9 | 7.61 |
| Cattle:Horse | Obsoletus complex | Europe | 0.96 | 37.7 | 48.8 | 0.77 |

## 4 Discussion

Our results indicate a complex pattern of host preference among different *Culicoides* species, reflecting both species-specific and regional variations in host selection. This systematic review revealed that while some *Culicoides* species demonstrated a strong preference for specific host types, others appeared more opportunistic, suggesting that host availability might play a significant role in their feeding behaviour. Awareness of this variability is crucial for understanding the dynamics of disease transmission and vector management.

The data collected in this review provide evidence of association in space and time between the insects and susceptible hosts, supporting their potential roles as vectors of pathogens and demonstrating one of the WHO criteria for vector recognition [1]. For some of the pathogens considered in this study not all of the other criteria are met. For example, transmission to hosts from infectious vectors has not been demonstrated for bovine ephemeral fever virus [190, 191].

Studies have shown that sensory structures of *Culicoides* species (palps and antennae) can determine whether they prefer feeding on birds or mammals [192, 193] or are opportunistic feeders [194]. Studies have also found evidence of *Culicoides* feeding on other insects including common fruit flies (*Drosophila melanogaster*) [21] and *Aedes*, *Culex* and *Anopheles* mosquitoes [195]. If *Culicoides* are regularly feeding on other insects which feed on the blood of vertebrate hosts, for example mosquitoes, the origin of the blood meals present may reflect the foraging behaviours of the mosquitoes, rather than of the *Culicoides*.

Our results mostly agree with Martínez-de la Puente et al. [7], who reviewed literature to find that most European *Culicoides* species feed on multiple vertebrate species, but present clear preferences for birds or mammals. However, our results for the Obsoletus group found the mean percentage of blood meals that came from birds in locations where birds were present was 30%, with a maximum of 50% in one study. This contradicts results from Martínez-de la Puente et al. [7] which suggested that all members of the Obsoletus group and complex were mammalphilic. Hopken et al. [38] found that many American *Culicoides* species were more opportunistic feeders than was thought prior to the study which identified blood meals of *Culicoides* caught in North America.

In our blood meal review we included data points for all competent *Culicoides* species and susceptible hosts at a location. This approach considers that in different settings *Culicoides* may feed on different hosts. Previous studies have suggested that the obsoletus complex prefers cattle to horses [184], however our review found that where horses are present, horse blood constitutes the highest percentage of the blood meals. These results do not necessarily contradict each other, but show the importance of considering the other host species in a setting and how the impact of this may vary for each species.

This variation in the origin of blood meals for some species was also observed in the host preference review. While similar results were found when comparing the number of a *Culicoides* species caught on or nearby some host pairs, these were often not consistent between studies, including studies at the same location. Although trends observed in the blood meal and host preference reviews generally agreed, this was not always the case. This highlights the need for a deeper understanding of the interaction between host preference and host availability for opportunistic feeders and its possible impact on disease transmission.

Host preference has been shown to be inherited for *Aedes* [196] and *Anopheles* [197] mosquitoes, with offspring more likely to select the same host as generations before them. If this trend is also true for *Culicoides* then this could contribute to the variations in results found in the host preference. If some hosts are usually less present at the location of the trial, the host preferences of local vectors could be more skewed towards host species that are more abundant in the area. More investigation into the innate host preferences of *Culicoides* spp. is required, as well as knowledge of the host distributions around trial sites to verify the reliability of these results.

The number of *Culicoides* collected from light traps near hosts is less than the number collected by aspiration directly from the hosts [13, 198]. Elbers and Gonzales [13] showed that the number of *Culicoides* caught by net sweeping significantly declined with increasing distance from hosts. In contrast, the number of *Culicoides* caught increased with increasing distance of hosts from the light trap.

Other viruses than those reported have been associated with *Culicoides* (including Ingwavuma virus, Jatobal virus, Manzanilla virus, Iquitos virus, Madre de Dios virus, Pintupo virus, Sango virus and Sathuperi virus [70]). However transmission has not been linked to specific species, often due to testing pooled *Culicoides* species.

## 5 Conclusion

Our findings suggest that future research should explore the drivers of host selection in more detail. We recommend the data on host selection needs to be assessed for a range of settings and at different locations in order to understand the foraging behaviours of *Culicoides* species. When using these data to parameterise mathematical models for disease transmission, the influence on host selection due to the variation in the settings in which hosts are found should be considered. Parameterising a model using data from one study is likely unreliable. Additionally, blood meal analysis with the distributions of host species usually found at a location are potentially more reliable than host preference studies comparing the number of *Culicoides* found on or near a host if one of the hosts is not usually present at the location.

These insights are instrumental for vector control strategies. Understanding that some vectors have fixed preferences helps in targeting control measures more effectively, focusing on the hosts that are most likely to be bitten in a particular setting. However, the adaptability in host preference observed in some species also warns against a one-size-fits-all approach to vector management. It emphasises the need for regional assessments of vector behaviour and host interactions to tailor interventions appropriately.

## Data availability

Supplementary file 1 contains all data gathered during the blood-meal systematic search. Supplementary file 2 contains summary statistics for the blood meal review for all *Culicoides* species and corresponding susceptible hosts. Supplementary file 3 contains results for all host pairs investigated for each *Culicoides* species in the host-preference study.

## Ethical approval

The authors confirm that the ethical policies of the journal, as noted on the journal’s author guidelines page, have been adhered to. No ethical approval was required.

## Author contributions

**ELF:** Conceptualisation, Methodology, Software, Programming, Validation, Formal analysis, Investigation, Data curation, Writing - Original Draft, Writing - Review & Editing, Visualisation, Project administration, Funding acquisition; **MJT:** Methodology, Writing - Review & Editing, Visualisation, Project administration, Funding acquisition; **JMD:** Writing - Original Draft, Writing - Review & Editing, Visualisation, Funding acquisition

